# From Patient to Tumor Organoid: Culture Protocol Choice Controls Glioblastoma Tumor Architecture and Identity

**DOI:** 10.64898/2026.04.28.721493

**Authors:** Jana Slovackova, Ondrej Bernatik, Katarina Cimborova, Martin Barak, Michal Hendrych, Karolina Kocourkova, Marie Sulcova, Jaroslav Olha, Katerina Amruz Cerna, Zdenek Hodny, Radim Jancalek, Dasa Bohaciakova

## Abstract

**Background:** Patient-derived tumor organoids are widely used in cancer research, yet the biological impact of tissue processing during model generation remains unclear. Fragment-based and dissociation-based approaches are commonly assumed to trade fidelity for uniformity, but their molecular consequences remain incompletely defined.

**Methods:** We performed a proteome-wide comparison of fragment-based (CUT) and dissociation-based (DIS) glioblastoma organoid protocols using quantitative mass spectrometry. Organoids from multiple patient tumors were cultured under growth factor–free or growth factor–supplemented conditions and compared with matched primary tissue.

**Results:** Both protocols produced technically robust glioblastoma organoids when maintained in their native media. However, CUT organoids matched the reproducibility of DIS cultures while preserving a broader extracellular matrix repertoire and networks linked to collagen assembly, vascular support, and cell–matrix signaling. DIS cultures were biased toward exogenous basement membrane components and proliferative, growth factor–responsive states. Across tumors, CUT organoids consistently showed greater proteomic similarity to matched primary tissue, retaining neural, glial, stromal, and extracellular features largely absent from DIS models.

**Conclusions:** Fragment-based glioblastoma organoids can be both reproducible and biologically faithful. Tissue dissociation acts as a major perturbation that reshapes extracellular matrix organization, cellular states, and tumor identity, making protocol choice a critical determinant of model fidelity and translational relevance.

## Introduction

Three-dimensional tumor organoids have become widely used models for studying human cancer biology, offering advantages over two-dimensional cultures by preserving aspects of tumor architecture, cellular heterogeneity, and intercellular signaling. In glioblastoma, IDH-wildtype (glioblastoma), patient-derived organoids are increasingly applied to investigate tumor-intrinsic programs, therapeutic responses, and microenvironmental interactions(1–3). However, organoids are generated using diverse culture protocols, and it remains unclear how methodological choices influence their molecular composition and fidelity to the parental tumor.

Two major strategies dominate tumor organoid culture. Fragment-based, or “cut-and-culture” (CUT), approaches maintain small pieces of tumor tissue without complete dissociation, preserving native cell–cell contacts, extracellular matrix (ECM), and stromal components(2). In contrast, dissociation-based (DIS) protocols enzymatically digest tumors into single cells that are reaggregated and cultured in growth factor– supplemented media, often embedded in Matrigel(1). Dissociation-based cultures are generally considered more uniform and technically reproducible, whereas fragment-based systems are often viewed as intrinsically heterogeneous(4,5). Whether this perceived heterogeneity reflects technical variability or instead retention of biologically meaningful tumor complexity remains unresolved.

Despite the rapid adoption of glioblastoma organoid models, direct comparisons of how these systems are generated remain surprisingly rare, with most studies centered on a single protocol. As a result, model performance has been judged primarily by morphology, immunohistochemistry, or transcriptomic profiling of the chosen culture method (1,6,7). While informative, these approaches do not directly capture protein-level regulation or extracellular matrix composition, which play central roles in tumor behavior. The ECM is not merely structural but actively regulates invasion, angiogenesis, mechanotransduction, and therapy resistance(8–11). Enzymatic dissociation disrupts endogenous ECM networks(12) and the loss of cell– matrix interactions, combined with enzymatic modification of surface receptors, may itself contribute to cellular reprogramming(13,14). And while exogenous matrices such as Matrigel provide a simplified, compositionally fixed substitute(15), they do not fully recapitulate patient-specific tumor microenvironments(16). How different organoid protocols preserve or replace native tumor ECM has not been systematically examined.

Mass spectrometry–based proteomics offers a powerful approach to address these questions by directly quantifying functional effectors, including ECM and secreted proteins that are often poorly represented at the transcript level(17,18). Although proteomic profiling has been extensively applied to glioblastoma tissues and in vitro models, including primary tumors, stem cell–derived cultures, and 2D/3D cell line systems(19–22), its application to patient-derived glioblastoma organoids remains limited. Existing studies have either focused on indirect readouts, such as extracellular vesicle proteomics from organoid cultures(23), short-term 3D cultures derived from dissociated glioblastoma stem cells(22), or targeted signaling analyses using antibody-based approaches(7). While these reports demonstrate the feasibility of proteomic interrogation in glioblastoma 3D systems, they do not provide a comprehensive, proteome-wide comparison of distinct organoid derivation strategies, nor do they address long-term organoid cultures that preserve tumor architecture.

Here, we present a systematic proteomic comparison of fragment-based and dissociation-based glioblastoma organoid cultures. By decoupling the effects of tissue processing and culture medium, quantifying technical and biological variability, and directly comparing organoids with matched primary tumors, we show that fragment-based organoids are technically robust and preserve endogenous ECM networks, cellular diversity, and tumor-specific proteomic features more faithfully than dissociation-based systems. These findings provide a molecular framework for selecting organoid culture strategies based on experimental goals and highlight the importance of proteomic benchmarking in tumor organoid research.

## Methods

### Primary tumor collection

Primary human tumor tissue was obtained from six patients undergoing surgical resection, with written informed consent and in accordance with institutional ethical approval (protocol number EK-FNUSA-27/2023). Clinical and pathological data are summarized in Supplementary **Table 1**. Fresh tissue (1-2 cm^3^) from a single resection site was transported from the operating room to the laboratory in ice-cold Hibernate-A medium (Gibco) supplemented with 1x B27 (Gibco) and 1x GlutaMAX (Gibco) and processed within 1– 2 hours (hrs) of resection. Viable tumor fragments from the tumor core were retained for organoid culture and proteomic analysis. A fraction of each tumor was snap-frozen in liquid nitrogen or fixed in 4% formaldehyde for histological and reference analyses.

### Fragment-based (architecture-preserving) organoid culture

Tumor fragments were processed following previously described fragment-based organoid culture protocols(2) with minor modifications. This protocol is here referred to as CUT protocol. Briefly, tissue was cut into approximately 0.5–1 mm^2^ fragments using sterile scalpels. Fragments were gently washed in Hibernate-A medium (Gibco) supplemented with 1xpenicillin–streptomycin (Gibco) and 1xAmphotericin B (Gibco). Pieces containing substantial amounts of necrosis or surrounding brain tissue were removed and only macroscopically viable fragments with visible tumor cell density were selected for culture. Because this protocol preserves native tissue architecture, the exact starting cell number per fragment was not determined. The remaining tumor pieces were grown in GBO medium lacking exogenous growth factors, consisting of 50% DMEM:F12 (Gibco), 50% Neurobasal (Gibco), 1x GlutaMAX (Gibco), 1x NEAAs (Gibco), 0.5x Zell Shield (Minerva), 1x N2 supplement (Gibco), 1x B27 w/o vitamin A supplement (Gibco), 1x 2-mercaptoethanol (Gibco), and 2.5 μg/ml human insulin (Sigma-Aldrich). No Matrigel was used in this culture. Medium was changed every 2–3 days, and cultures were maintained on an orbital shaker at 90 rpm at 37 °C, 5% CO_2_, and 95% relative humidity. Organoids were monitored by phase-contrast microscopy and collected for downstream analyses after 30 days in culture.

### Dissociation-based (single-cell reaggregation) organoid culture

For dissociation-based cultures (here referred to as DIS protocol), tumor tissue was minced and enzymatically digested according to Stemcell technology protocol(24) using papain (Wortinghton) and DNase I (StemCell Technology) solution for 60 minutes, shaking at 90 rpm at 37 °C. The resulting suspension was filtered through a 37-µm strainer (PluriSelect) and centrifuged at 300xg for 5 minutes. Red blood cells were removed using RBC lysis buffer (PluriSelect) if necessary. Viable cells were counted by trypan blue exclusion. Cells were then resuspended in cold growth-factor-reduced Matrigel (Corning) and plated as 5 µL droplets. A defined number of viable cells was used to establish each Matrigel droplet, corresponding to approximately 20,000 cells per 5 µL droplet. Solidified droplets were covered with NBM medium containing the following components: Neurobasal medium (Gibco) supplemented with recombinant human EGF (20 ng/mL; Peprotech) and recombinant human FGF2 (20 ng/mL; Peprotech), 1x B27 (Gibco), 1x GlutaMAX (Gibco), 1x Sodium pyruvate (Gibco), and 0.5x Zell Shield (Minerva). Medium was replaced every 2–3 days. Dissociation-derived spheroids were cultured for 30 days prior to collection and reached a similar size to those cultured by CUT procol.

### AV-GFP Labeling of tumoroids

To visualize tumor cells within organoids, cultures were labeled with adenoviral GFP (AV-GFP, Vector Biolabs). Tumor organoids of 1mm^2^ in average size were incubated with AV-GFP in complete culture medium at 1x 10^7^ PFU to allow viral penetration and labeling of cells. Organoids were exposed to the virus for 24 hours at 37 °C and 5% CO_2_ under gentle agitation. After incubation, organoids were washed twice with fresh medium to remove residual virus, then maintained under standard culture conditions. GFP expression was assessed by fluorescence microscopy to confirm successful labeling and to monitor organoid structure and cell distribution over time. Nuclei were counterstained with 4′,6-diamidino-2-phenylindole (DAPI). The same procedure was used for the CUT and DIS protocols.

### Medium comparison experiments

To distinguish protocol-dependent effects from medium-dependent effects, both organoid derivation methods were cultured in parallel under two media conditions. CUT organoids were grown either in their native GBO medium or in NBM medium, whereas DIS organoids were grown either in their native NBM medium or in GBO medium. All conditions were established as matched technical replicates and processed in parallel.

### Histological analysis and immunohistochemistry

Organoids and primary tumor tissue were fixed in 4% formaldehyde for 24 hrs at 4 °C, washed in PBS, dehydrated through graded ethanol, and embedded in 3% low-gelling Agarose (Sigma) before paraffin-embedding. Sections (2 µm) were cut using a rotary microtome and mounted on positively charged slides (TOMO adhesion slides, Matsunami). For routine morphology, sections were stained with hematoxylin and eosin (H&E) following standard procedures.

Immunohistochemistry (IHC) was performed using an IHC/ISH Slide Staining System (BenchMark ULTRA, Ventana Medical Systems) according to the manufacturer’s manual. Sections stained with primary antibodies against: GFAP, S100, NSE, and CD31 (antibody details and dilutions listed in **Supplementary Table 2**). After washing, visualization was performed with the Ultra View DAB IHC Detection Kit (Ventana Medical Systems). The specimens were then counterstained with hematoxylin Gill No. 3 (Sigma-Aldrich) for 1 minute, blued in tap water, dehydrated, and coverslipped. Images were acquired using a ZEISS Axioscan 7 microscope (Zeiss, Oberkochen, Germany). No image manipulations that could affect data interpretation were performed.

### Proteomics

Proteins from samples were extracted in SDT buffer (4% SDS, 0.1M DTT, 0.1M Tris/HCl, pH 7.6) in a thermomixer (Eppendorf ThermoMixer C, 60 min, 95°C, 750 rpm). Subsequently, all samples were centrifuged (15 min, 20,000 x g), and the supernatants (ca. 50 μg of total protein) were used for filter-aided sample preparation (FASP) as described before(25) using trypsin as the protease (enzyme-to-protein ratio of 1:50; sequencing grade; Sigma-Aldrich). The resulting peptides were extracted into an LC-MS vial by 2.5% formic acid (FA) in 50% acetonitrile (ACN) and 100% ACN with the addition of polyethylene glycol (final concentration 0.001%)(26) and concentrated in a SpeedVac concentrator (Thermo Fisher Scientific).

LC-MS/MS analyses of all peptides were done using the UltiMate 3000 RSLCnano system (Thermo Fisher Scientific) connected to the timsTOF HT mass spectrometer (Bruker). Before LC separation, tryptic digests were online concentrated and desalted using a trapping column (300 μm × 5 mm, μPrecolumn, 5 μm particles, PepMap™ Neo Trap Cartridge, Thermo Fisher Scientific). The trap column was then washed with 0.1% trifluoroacetic acid and the peptides were eluted in backflush mode from the trapping column onto an analytical column (Aurora C18, 75μm ID, 250 mm long, 1.7 μm particles, PN AUR3-25075C18-CSI; Ion Opticks) by linear 90m gradient program (flow rate 200 nl.min^-1^, 3-42% of mobile phase B; mobile phase A: 0.1% FA in water; mobile phase B: 0.1% FA in 80% ACN) followed by a system wash using 80% of mobile phase B. Equilibration of the trapping column and the analytical column was done before sample injection to sample loop. The analytical column was installed in the Captive Spray ion source (Bruker; temperatures set to 50ºC) according to the manufacturer’s instructions. Spray voltage was set to 1.5kV.

MS data were acquired in data-independent acquisition (DIA) mode with m/z range of 100-1700 and 1/k0 range of 0.6-1.4 V×s×cm^-2^. DIAparameters.txt file defined 21 Th windows in m/z 400-1000 precursor range using two steps for each PASEF scan and cycle time of 100ms locked to 100% duty cycle.

DIA data were processed in DIA-NN (version 2.1.0)(27,28) in library-free mode against the modified UniProtKB protein database (https://www.uniprot.org/proteomes/UP000005640) for *Homo sapiens*; version 2025/04, number of protein sequences: 20,663. The original UniProtKB database was supplemented by iRT peptide sequences (Biognosys). No optional, but carbamidomethylation as fixed modification and trypsin/P enzyme with 1 allowed missed cleavages and peptide length 7-30 were set during the library preparation. False discovery rate (FDR) control was set to 1% FDR. MS1 and MS2 accuracies as well as scan window parameters were set based on the initial test searches (median value from all samples ascertained parameter values). MBR was switched on.

### Data Analysis

Downstream analysis was performed in R(29) (version 4.4.1). Protein groups annotated as contaminants or identified on less than 2 proteotypic peptides were excluded. Proteins were separated into two groups, quantitatively (found in at least 3 samples in all sample groups and in at least 4 samples in two sample groups) and qualitatively (found in 4 or more samples in one sample group, while not found in more than 2 samples in other sample groups). Proteins never detected in more than 3 samples in a sample group were excluded from the analysis. Quantitatively changed protein LFQ intensities were log2-transformed, and missing values were imputed using a left-shifted Gaussian distribution and used to compute global sample characteristics, like principal component analysis and clustering heatmaps. Principal component analysis (PCA), correlation matrices, and hierarchical clustering were used to assess sample relationships. Differential protein abundance was calculated using limma or linear mixed-effects models, with multiple-testing correction by Adjusted p. value. Proteins with p. value < 0.05 and log2 fold-change ≥ 1 or ≤ -1 threshold were considered significantly changed. Functional enrichment analyses were performed using clusterProfiler_(4.14.6)(30,31) using GO(32–34), KEGG(35) and Reactome databases(36) and AnnotationDbi_(1.68.0), and org.Hs.eg.db (3.20.0) when necessary for annotations. Data visualization was performed using ggplot2 (4.0.0)(37), Pheatmap (1.0.13)(38), and UpSetR (1.4.0)(39).

Matrisome annotations were retrieved from the MatrisomeDB(40) (MatrisomeAnalyzeR, 1.0.1) resource for ECM-focused analyses. Lists of matrisome proteins enriched in CUT/GBO and DIS/NBM samples were submitted to STRING (version 12.0)(41,42) using human gene symbols as input identifiers (organism: *Homo sapiens*). STRING networks were generated using the full STRING network with a minimum required interaction score of 0.4 (medium confidence) and the following interaction sources enabled: experiments, curated databases, co-expression, gene neighborhood, gene fusion, and co-occurrence. No additional interactors were added (first and second shell = none), disconnected nodes were hidden, and network clustering was performed using the Markov Cluster Algorithm (MCL) with an inflation parameter of 1.5.

## Results

### 1. Proteomic study design to compare two commonly used tumor organoid protocols

To systematically assess how culture methodology influences the molecular composition of patient-derived tumor organoids, we focused on two widely used protocols that differ fundamentally in how they process tumor tissue(1,2). Both approaches were implemented according to published procedures and subsequently optimized in our laboratory. As schematized in **Fig. 1A**, the CUT protocol involves mechanically sectioning tumor tissue into small fragments, which are cultured directly in GBO medium without exogenous growth factors or Matrigel, as originally described by Jacob and colleagues(2). This strategy is intended to preserve native tissue architecture and stromal elements while allowing the tumor to expand in a three-dimensional context. Representative brightfield images, H&E-stained sections, and AV-GFP–labeled cultures are shown in **Fig. 1B**.

**Figure 1.**
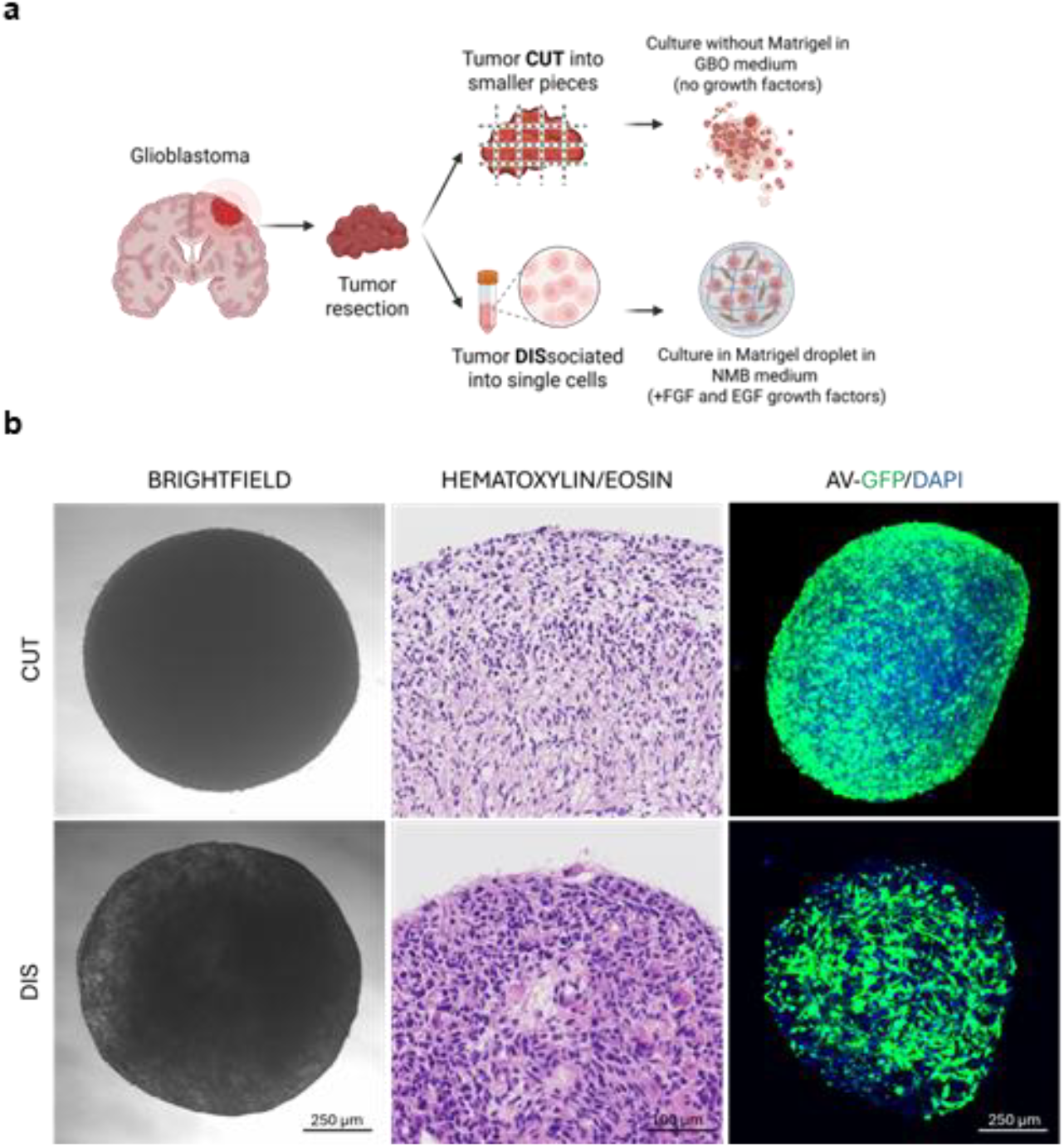
Study design for proteomic comparison of two commonly used tumor organoid protocols. **(A)** Schematic overview of experimental workflow. Primary human tumor tissue was processed using two established organoid culture strategies: the CUT protocol, in which tumor fragments are mechanically sectioned and maintained in GBO medium lacking exogenous growth factors, and the DIS protocol, in which tissue is enzymatically dissociated into single cells and replated in Matrigel droplets in growth factor–supplemented NBM medium (containing EGF and FGF). Experimental arms included (i) evaluation of technical variability, (ii) comparison of the effects of culture medium, and (iii) assessment of biological variability across independent tumors. **(B)** Brightfield images of organoids cultured for 30 days, histological sections stained with hematoxylin and eosin, and AV-GFP labeled cells in organoids from individual culture conditions. Nuclei were counterstained with DAPI.

In contrast, the DIS protocol(1) relies on enzymatic digestion of tumor tissue into single cells, followed by reaggregation in Matrigel droplets and maintenance in growth factor–supplemented medium containing EGF and FGF2. This approach is believed to yield more uniform spheroid-like structures (**Fig. 1B**) and is considered technically more reproducible. Because both protocols(1,2) are widely used and are associated with distinct, and sometimes conflicting, assumptions about how faithfully they recapitulate primary tumors, we designed a comparative proteomic study to evaluate their molecular consequences in a controlled and systematic manner. In the following sections, we examine technical robustness, the influence of culture medium, biological variability across tumors, and the extent to which each protocol resembles the matched primary tumor.

### 2. CUT and DIS protocols generate comparably homogeneous organoids under native conditions

To first evaluate the technical robustness of the two organoid culture protocols and assess the degree of similarity among tumor organoids generated by each method, we first performed a systematic proteomic comparison of technical replicates across culture media (**Fig. 2A** and **Supplementary Table 3**). Organoids were generated from a single patient sample using either the DIS or CUT protocol (10 technical replicates per protocol). For each protocol, organoids were maintained for 30 days in two types of media: one containing growth factors (NBM) and one without growth factors (GBO), yielding a total of 20 cultured samples. In addition, the corresponding primary (non-cultured) tumor tissue was included as a reference. Interestingly, correlation analysis of proteomics results revealed that, when cultured in their “native” media (i.e., DIS organoids in NBM and CUT organoids in GBO), both protocols produced highly consistent proteomic profiles across technical replicates, with clear clustering by condition and minimal dispersion (**Fig. 2B**). CUT-derived tumor organoids also remained comparatively similar when cultured in NBM medium, suggesting that CUT is relatively insensitive to the presence of growth factors and other “non-native” media supplements. In contrast, DIS-derived organoids cultured in GBO medium deviated markedly from all other DIS samples, indicating that the absence of growth factors, and possibly other GBO media supplements, substantially alters the proteomic composition of DIS organoids. These trends were further supported by principal component analysis, in which DIS organoids grown in GBO medium formed a distinct and separated cluster (**Fig. 2C**).

**Figure 2.**
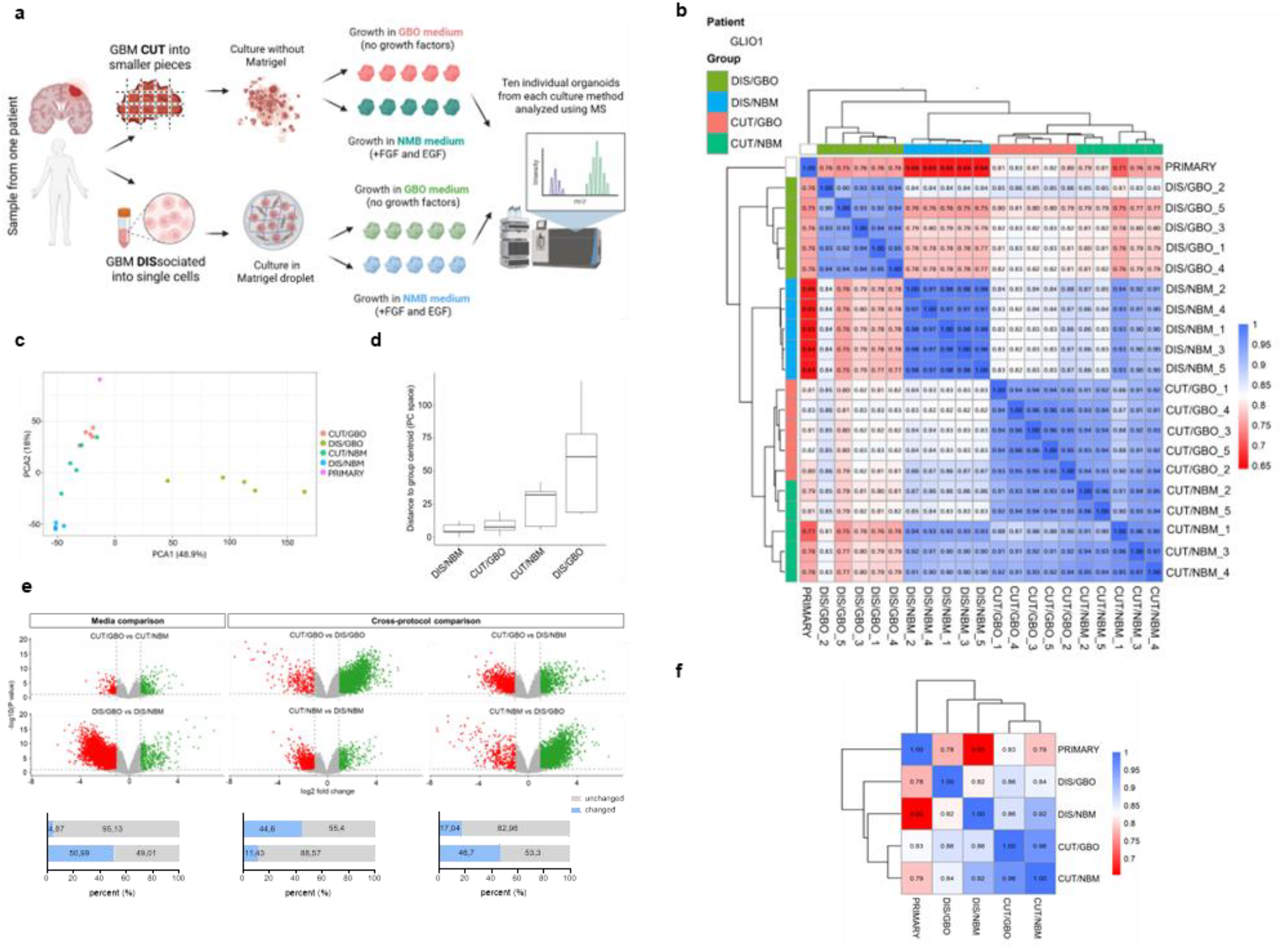
CUT and DIS protocols generate comparably homogeneous organoids under native conditions. **(A)** Experimental design. Organoids were generated from a single patient tumor using either CUT or DIS based protocols (10 technical replicates each) and cultured for 30 days in growth factor–free GBO medium (5 technical replicates) or growth factor–containing NBM medium (5 technical replicates). The matched primary tumor was included as a reference; **(B)** Pearson’s correlation heatmap of proteomic profiles; **(C)** Principal component analysis (PCA); **(D)** Distance-to-centroid analysis of individual organoid samples within each culture condition, calculated from proteomic profiles. Shorter distances indicate lower variability and higher technical consistency within a group; **(E)** Volcano plots and pairwise similarity analysis across conditions; **(F)** Pearson’s correlation to the primary tumor.

Furthermore, to quantitatively assess variability within each condition, we next evaluated the dispersion of individual samples by calculating their distance to the respective group centroid (**Fig. 2D**). As expected, DIS organoids cultured in their native NBM medium exhibited very low variability across technical replicates. Importantly, CUT organoids cultured in GBO medium displayed a similarly low degree of variability, comparable to that observed for DIS/NBM organoids. In contrast, increased variability was observed when CUT organoids were cultured in NBM medium, and the highest level of dispersion was detected in DIS organoids maintained in GBO medium. Together, these analyses demonstrate that while the DIS protocol is strongly dependent on its native medium to maintain proteomic stability, both protocols produce relatively homogeneous tumor organoids when cultured under their respective optimized conditions.

To better understand how protocol and medium shape organoid composition, we next quantified proteome similarity across conditions (**Fig. 2E**). CUT organoids cultured in GBO versus NBM media were highly consistent, sharing 96.51% of proteins with similar intensity. In contrast, DIS organoids showed only 52.17% similarity when grown in GBO medium rather than their native NBM medium, indicating marked sensitivity to the withdrawal of growth factors and other media components. Cross-protocol comparisons further supported this pattern. DIS organoids maintained in GBO medium shared only 58.90% similarity with CUT/GBO cultures and 57.04% similarity with CUT/NBM cultures. By comparison, DIS organoids grown in their native NBM medium were much more aligned: they shared 89.13% similarity with CUT/NBM and 82.79% similarity with CUT/GBO. Taken together, these analyses show that both protocols generate broadly comparable organoids when cultured under their optimal conditions, whereas placing DIS organoids in GBO medium substantially reduces proteomic similarity.

Finally, we compared all cultured organoids to the matched primary tumor sample (**Fig. 2F**). CUT organoids grown in GBO medium exhibited the highest correlation to the primary tissue, regardless of the medium. DIS organoids cultured in NBM medium showed a moderate degree of deviation, whereas DIS organoids maintained in GBO medium displayed the largest proteomic shift from the primary tumor. These findings highlight that both protocol choice and medium composition critically influence the extent to which tumor organoids recapitulate the primary tumor proteome.

### 3. Biological variability reveals distinct molecular signatures between CUT and DIS organoids

To further evaluate how culture methodology shapes tumor organoid biology across independent tumors, we next analyzed tumor resections from five individual patients. For each patient, 12 organoids were generated: three cultured as CUT/GBO, three as CUT/NBM, three as DIS/GBO, and three as DIS/NBM. For proteomic analysis, organoids grown under the same conditions were pooled, yielding one sample per condition. This resulted in four organoid samples per patient. Together with matched primary tumor sample per each patient, we generated a total of 25 samples for proteomic analysis (**Fig. 3A**). PCA and Pearson’s correlation analysis demonstrated that DIS protocol-generated organoids maintained in GBO medium again underperformed and were distinct from the remaining samples (**Fig. S1A, B**), further supporting that non-native medium is poorly tolerated in the DIS system. In contrast, CUT-derived organoids exhibited relatively tighter clustering across donors, suggesting greater biological consistency.

**Figure 3:**
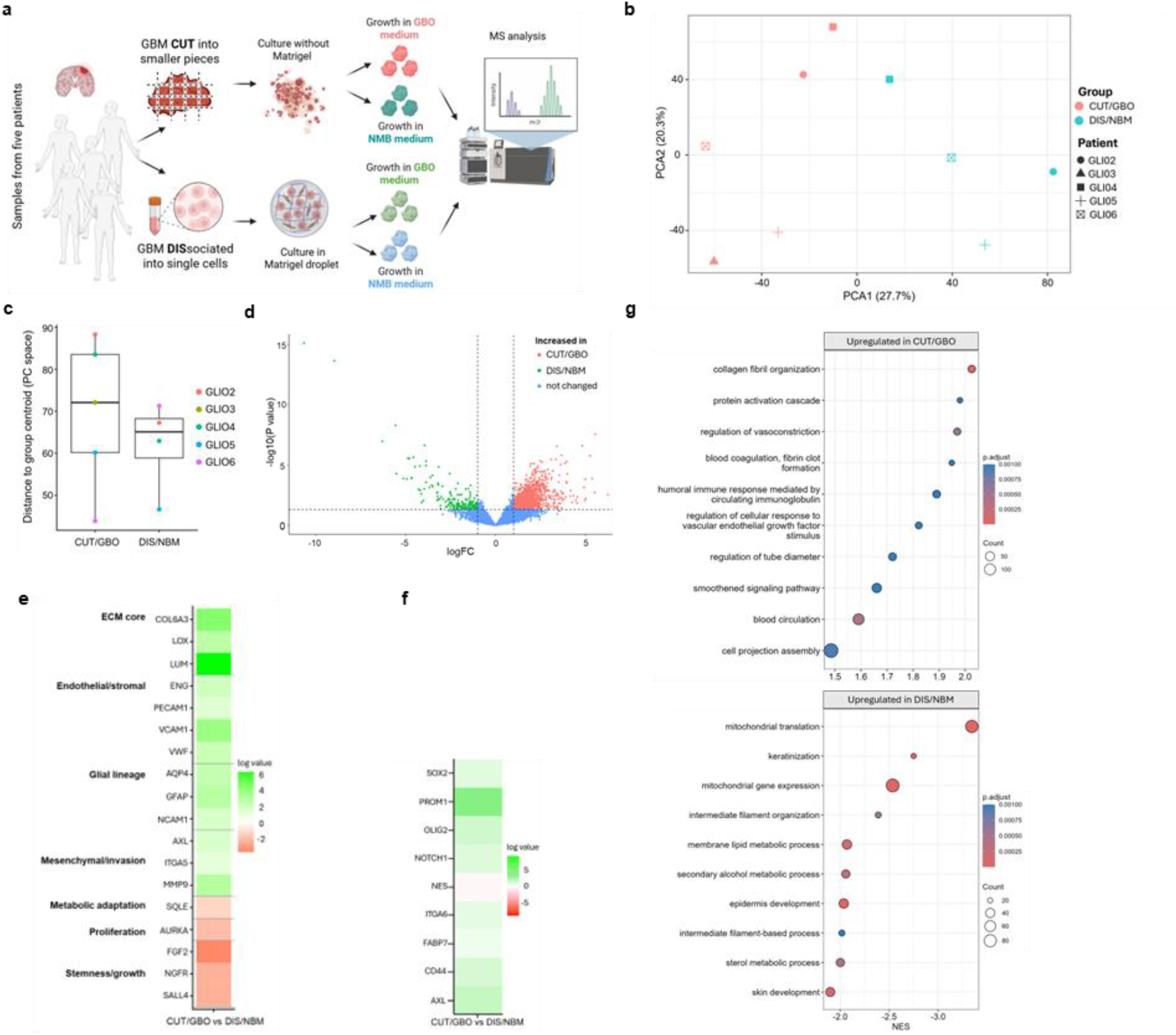
Biological variability and protocol-specific molecular signatures in patient-derived glioblastoma organoids. **(A)** Experimental design for analysis of biological variability across five independent patient tumors, including CUT and DIS organoids cultured in GBO or NBM media and matched primary tumor samples. **(B)** PCA of CUT/GBO and DIS/NBM organoids across biological replicates. **(C)** Quantification of variance across biological replicates cultured under native conditions. **(D)** Volcano plot showing differential protein abundance between CUT/GBO and DIS/NBM organoids across tumors. **(E)** Enrichment analysis of proteins differentially abundant between CUT/GBO and DIS/NBM organoids. **(F)** Targeted heatmap displaying representative proteins associated with enriched biological processes and additional significantly regulated proteins. **(G)** Relative abundance of selected glioblastoma lineage- and stemness-associated marker proteins across CUT/GBO and DIS/NBM organoids.

Therefore, to further quantify shared and unique proteomic features between the culture strategies, we compared organoids generated under two selected conditions in their respective native media: the CUT protocol in GBO medium and the DIS protocol in NBM medium. These conditions were selected because they represent the canonical use of each protocol and performed best in our analyses of technical and biological variability. We thus next performed principal component analysis restricted to biological replicates cultured under these native conditions (**Fig. 3B**) and quantified variance across biological samples (**Fig. 3C**). In both protocols, biological variability between patient-derived samples was substantial and clearly exceeded technical variability. Importantly, this high degree of biological heterogeneity was observed irrespective of the culture strategy and was comparable between CUT and DIS organoids, with a tendency toward slightly greater variance in DIS-derived samples. These results indicate that inter-patient biological differences dominate proteomic variability, while protocol-dependent effects represent a secondary, though structured, source of variation. Processed data are available in **Supplementary Table 4**.

Direct differential abundance analysis across tumors confirmed broad differences between CUT/GBO and DIS/NBM organoids as shown by a volcano plot (**Fig. 3D**), with more proteins being significantly overrepresented in the CUT protocol compared to the DIS protocol. Unbiased enrichment analysis of proteins differentially abundant between CUT/GBO and DIS/NBM organoids identified extracellular matrix organization, collagen assembly, cell adhesion, and cilium organization as the most significantly enriched biological processes in CUT/GBO cultures (**Fig. 3E**). In contrast, DIS/NBM organoids showed enrichment of biosynthetic and proliferation-associated pathways, including translational activity, and lipid/cholesterol metabolism, suggesting a growth factor–responsive and culture-adapted spheroid state. DIS/NBM cultures were also characterized by enrichment of basement membrane components, including laminins and type IV collagen, likely reflecting the intrinsic use of Matrigel.

To illustrate these findings, we visualized representative proteins contributing to the enriched biological processes, together with additional significantly regulated proteins functionally linked to these pathways, using a targeted heatmap **(Fig. 3F)**. Consistent with enrichment of extracellular matrix organization and collagen assembly, CUT/GBO organoids exhibited increased abundance of core ECM components (COL6A3, LOX, LUM), endothelial and stromal markers (ENG, PECAM1, VCAM1, VWF), and glial and neural lineage–associated proteins (AQP4, GFAP, NCAM1). CUT/GBO cultures were also enriched for mesenchymal and invasion-associated proteins, including AXL, ITGA5, and MMP9. In contrast, DIS/NBM organoids showed higher abundance of proteins linked to proliferation, stemness, and growth factor responsiveness, such as AURKA, FGF2, NGFR, and SALL4, as well as metabolic enzymes including SQLE. Notably, canonical glioblastoma stem cell markers, including SOX2, PROM1 (CD133), OLIG2, NOTCH1, ITGA6, FABP7, CD44, and AXL, were not selectively enriched in DIS cultures but were broadly retained in CUT/GBO organoids **(Fig.3G)**, with NES representing the only marker preferentially enriched in DIS/NBM. Together, these data demonstrate that fragment-based CUT organoids preserve extracellular matrix architecture, vascular and stromal features, and a broad spectrum of tumor cell identities, including stem-like populations. In contrast, dissociation-based cultures preferentially select for proliferative and growth factor–responsive states, underscoring that protocol choice imposes a strong and directional bias on both extracellular and cellular tumor programs.

### 4. CUT organoids preserve extracellular matrix organization that is largely lost in DIS spheroids

Given the pronounced differences in extracellular matrix–associated proteins observed between culture conditions, we next performed a focused analysis of the glioblastoma tumor organoid matrisome (**Fig. 4A**). This analysis assessed how protocol choice influences preservation of core matrisome proteins (collagens, ECM glycoproteins, and proteoglycans) and associated matrisome proteins (ECM regulators, ECM-affiliated proteins, and secreted factors).

**Figure 4.**
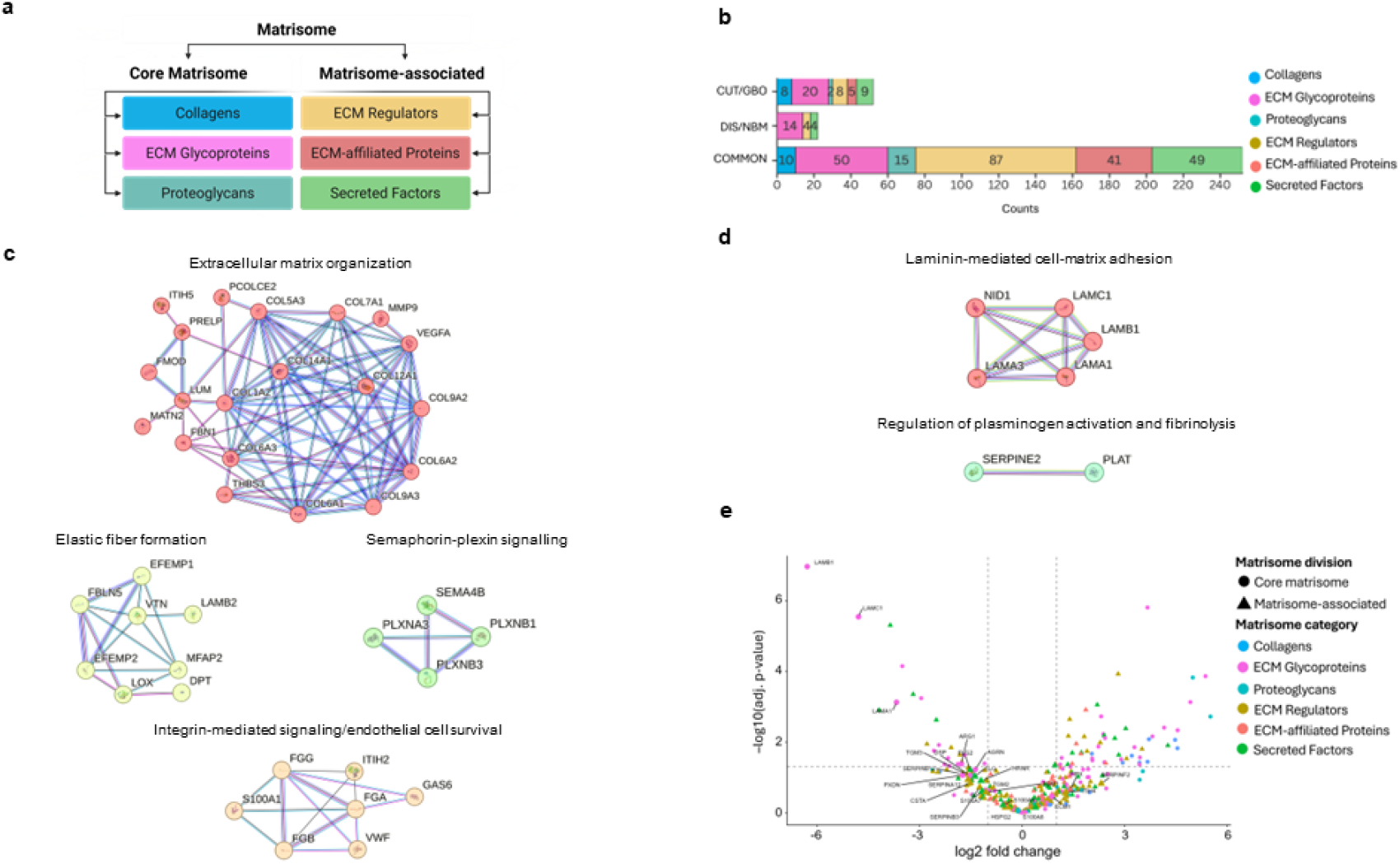
Fragment-based organoid culture preserves tumor extracellular matrix organization. **(A)** Schematic overview of core matrisome and associated matrisome protein categories; **(B)** Quantification of shared and uniquely detected matrisome proteins across major matrisome categories, including collagens, ECM glycoproteins, proteoglycans, ECM regulators, ECM-affiliated proteins, and secreted factors; **(C)** STRING protein–protein interaction network analysis of matrisome proteins enriched in CUT/GBO organoids; **(D)** STRING network analysis of matrisome proteins enriched in DIS/NBM organoids; **(E)** Volcano plot showing differential abundance of matrisome proteins between CUT/GBO and DIS/NBM organoids across tumors. Proteins abundant in Matrigel are highlighted.

Firstly, quantitative comparison of matrisome proteins between fragment-based CUT organoids cultured in GBO medium and dissociation-based DIS organoids cultured in NBM medium revealed that, across all categories, a substantial fraction of proteins was shared, indicating retention or de novo synthesis of a basic extracellular framework in both systems (**Fig. 4B**). However, in every matrisome category examined, CUT/GBO organoids consistently show proteins being enriched compared to the DIS/NBM. This difference was most pronounced for core matrisome components. Among collagens, 10 proteins were detected with similar intensity between conditions, whereas 8 additional collagens were enriched in CUT/GBO organoids and none were enriched in DIS/NBM. Similarly, for proteoglycans, 15 proteins were shared, 2 were enriched in CUT/GBO, and none were enriched in DIS/NBM. ECM glycoproteins represented the largest category overall, with 50 shared proteins; notably, CUT/GBO organoids were enriched for 20 ECM glycoproteins compared with 14 in DIS/NBM. Differences also extended to the associated matrisome. CUT/GBO organoids were enriched for twice as many unique ECM regulators (8 versus 4), while 5 ECM-affiliated proteins were detected enriched in CUT/GBO cultures. Likewise, CUT/GBO organoids exhibited more than twice the number of enriched secreted factors (9 versus 4). Together, these data demonstrate that although both protocols contain a shared baseline matrisome, CUT/GBO organoids retain a richer extracellular protein repertoire.

To assess how these differences translate into higher-order ECM organization, we performed STRING protein–protein interaction analysis on matrisome proteins enriched in each condition (**Fig. 4C**). CUT/GBO-enriched proteins formed multiple highly interconnected networks centered on extracellular matrix organization and collagen assembly, including collagens and key ECM structural and regulatory proteins such as LUM, FMOD, PRELP, FBN1, MMP9, and DPT. Additional networks were associated with elastic fiber formation (LOX, MFAP2, EFEMP2, FBLN5), integrin-mediated signaling and endothelial cell survival (FGG, FGB, FGA, ITIH2, VWF, GAS6, S100A1), and semaphorin–plexin signaling (SEMA4B, PLXNA3, PLXNB1, PLCB3), indicating coordinated preservation of stromal, vascular, and guidance-related ECM programs. In contrast, STRING analysis of DIS/NBM-enriched matrisome proteins revealed sparse and weakly connected networks (**Fig. 4D**), dominated by laminin-mediated cell–matrix adhesion (LAMA1, LAMA3, LAMB1, LAMC1, NID1) and a small fibrinolysis-related module (SERPINE2, PLAT), consistent with reliance on Matrigel-derived basement membrane components.

Finally, differential abundance analysis directly comparing matrisome proteins between CUT/GBO and DIS/NBM organoids further highlighted these differences (**Fig. 4E**). The volcano plot revealed a strong asymmetry, with the majority of significantly enriched matrisome proteins mapping to the CUT condition. In contrast, proteins preferentially abundant in DIS/NBM cultures were largely limited to basement membrane components known to be abundant in Matrigel, which were explicitly highlighted, including LAMB1, LAMC1, and LAMA1. This pattern mirrors the STRING network analysis, underscoring that CUT organoids preserve endogenous, tumor-derived ECM networks, whereas dissociation-based cultures likely predominantly reflect exogenous matrix inputs.

### 5. CUT organoids retain primary tumor features more faithfully than DIS cultures

Finally, to determine which culture strategy more closely recapitulates the molecular composition of the original tumor, we directly compared tumor organoids to their matched primary tumor samples. Principal component analysis, including both biological and technical replicate experiments, revealed a clear gradient of similarity (**Fig. 5A**). While some dispersion among samples was observed, CUT organoids consistently localized closer to primary tumor samples than DIS organoids. This pattern was reproducible across independent tumors and remained evident when technical replicates were included, indicating that the observed differences reflect biological rather than technical effects.

**Figure 5.**
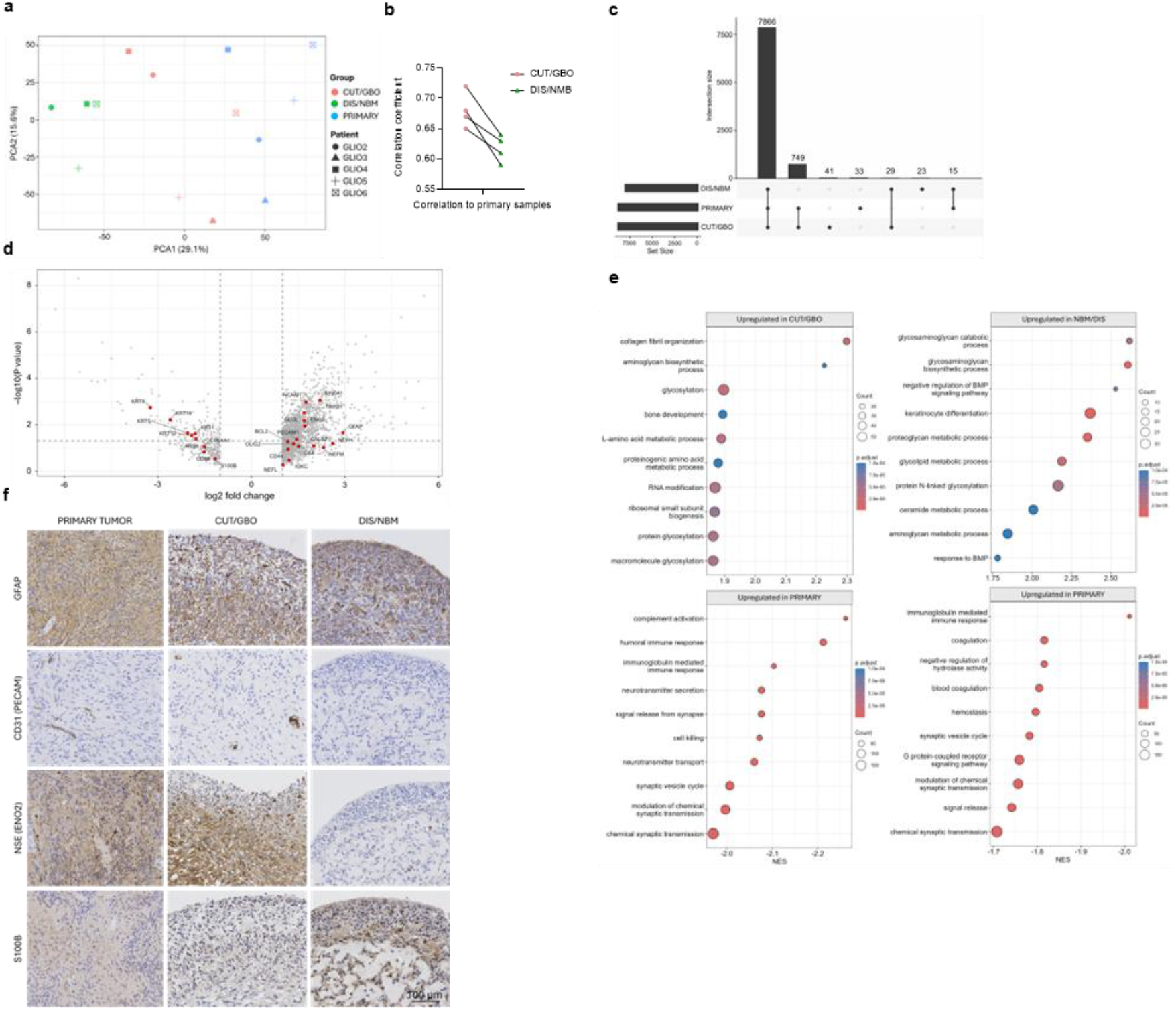
Comparison of tumor organoids with matched primary tumor samples. **(A)** PCA of proteomic profiles from primary tumor tissue, CUT-derived organoids, and DIS-derived organoids. Data include both biological replicates across multiple patients and technical replicates from a single patient. **(B)** Pairwise Pearson correlation coefficients between each organoid sample and its matched primary tumor, shown per culture condition. **(C)** UpSet plot illustrating the overlap of detected proteins among primary tumors, CUT organoids, and DIS organoids. Numbers indicate shared and condition-specific protein sets. **(D)** Volcano plot of differentially abundant proteins between CUT and DIS organoids relative to primary tumor samples, with selected proteins highlighted. **(E)** Representative histological and immunohistochemical staining of primary tumors and matched CUT- and DIS-derived organoids for selected markers. **(F)** Enrichment analysis of proteins retained in primary tumors or enriched in CUT or DIS organoids.

To quantitatively assess these relationships, we calculated pairwise correlation coefficients between each organoid sample and its corresponding primary tumor (**Fig. 5B**). Across all tumors, CUT organoids exhibited significantly higher correlation coefficients than their respective DIS organoids, indicating greater proteomic similarity to the parental tissue. In contrast, DIS-derived cultures showed consistently reduced similarity to primary samples, indicating more extensive proteomic remodeling following dissociation and reaggregation.

We next examined shared and condition-specific protein representation using an UpSet analysis (**Fig. 5C**). The majority of detected proteins (7,866) were shared across primary tumors, CUT organoids, and DIS organoids, reflecting preservation of core tumor programs in both systems. However, a substantial subset of proteins (749) was shared exclusively between primary tumors and CUT organoids, whereas only 15 proteins overlapped uniquely between primary tumors and DIS organoids. This striking imbalance indicates that CUT cultures retain a substantially larger fraction of primary-tumor–specific proteomic features.

Consistent with these observations, differential abundance analysis (**Fig. 5D**) showed that proteins enriched in CUT organoids relative to DIS included markers of glial and neural differentiation and tumor microenvironmental complexity, such as GFAP, PECAM1 (CD31), and ENO2 (NSE). In contrast, DIS organoids were enriched for proteins associated with simplified or stress-adapted states, including S100B, ARG1, COL4A1, and CD68, consistent with altered cellular composition and increased reliance on basement membrane–associated components. Histological and immunohistochemical analyses provided orthogonal validation of these findings (**Fig. 5E**). Staining for GFAP, PECAM1, and ENO2 showed closer spatial distribution and expression patterns between primary tumors and CUT-derived organoids than DIS-derived cultures, whereas S100B, enriched in DIS organoids, exhibited divergent patterns relative to the primary tissue.

Finally, we aimed to identify which primary tumor features are consistently lost during organoid derivation, irrespective of the protocol used. Enrichment analysis of proteins retained in primary tumors but depleted in both CUT and DIS organoids revealed strong enrichment for synaptic and immune-related processes (**Fig. 5F**), including synaptic vesicle cycling, neurotransmitter secretion, vesicle-mediated transport at synapses, complement activation, humoral immune responses, and B–cell–mediated immunity. In contrast, proteins enriched in CUT organoids were primarily associated with general cellular processes, including RNA modification, tRNA processing, ribosomal function, and glycosylation, whereas proteins enriched in DIS organoids were linked predominantly to non-specific metabolic pathways. Together, these analyses indicate that while both organoid systems lose specific neuronal and immune features present in primary tumors, fragment-based CUT organoids preserve a substantially broader and more faithful representation of the primary tumor proteome than dissociation-based cultures.

## Discussion

A central challenge in tumor organoid research and disease modeling in vitro is balancing technical compliance with biological reliability. Here, by systematically comparing fragment-based and dissociation-based glioblastoma organoid protocols using deep proteomic profiling, we demonstrate that protocol choice is not a neutral technical decision but a biologically consequential variable that shapes cellular states, extracellular matrix organization, and ultimately similarity to the parental tumor. Our data reveal that while both approaches generate viable, reproducible organoids, they differ in predictable, structured ways that have important implications for tumor modeling.

### Impact of tissue dissociation on cellular states and tumor identity

A prevailing assumption in the tumor organoid field is that dissociation-based protocols yield more homogeneous and reproducible cultures, whereas fragment-based approaches are intrinsically heterogeneous due to preservation of native tissue complexity and multicellular architecture(1,2,6,7). One of the key objectives of this study was, therefore, to directly test this assumption by quantitatively assessing the degree of heterogeneity in fragment-based CUT organoids relative to dissociation-based cultures at the proteomic level.

Interestingly, contrary to expectation, fragment-based CUT organoids cultured in their native medium displayed a high degree of proteomic consistency across technical replicates, comparable to that observed in dissociation-based organoids maintained under optimal conditions. This finding challenges the notion that CUT organoids are inherently heterogeneous and demonstrates that preservation of tissue architecture does not necessarily compromise technical robustness. Instead, the variability observed in CUT cultures largely reflects retained biological complexity rather than increased technical noise. These observations are particularly notable in light of prior studies showing that enzymatic dissociation itself can induce stress responses, metabolic reprogramming, and altered molecular states that may artificially increase apparent homogeneity while reshaping cellular identity(43,44).

Beyond technical variability, protocol choice imposed a consistent and directional shift in cellular-state composition. CUT organoids retained proteins associated with differentiated neural and glial programs, endothelial and stromal components, as well as cancer stem cells, consistent with preservation of multicellular tumor architecture. In contrast, dissociation-based cultures were biased toward growth-responsive, metabolically active states, as evidenced by enrichment for proliferation-associated regulators and lipid and cholesterol biosynthesis pathways. Importantly, these differences were reproducible across independent patient-derived tumors, indicating that dissociation acts as a structured biological perturbation rather than a source of random variability. Our findings thus align with transcriptomic studies demonstrating that tissue dissociation induces conserved stress and metabolic programs and can alter cellular identity(45,46), and extend these observations by showing that such effects persist at the proteomic level in long-term organoid cultures. In the context of glioblastoma, the combination of dissociation and sustained growth factor exposure favored expansion of progenitor-like cellular states, consistent with established growth factor dependencies of stem-like tumor populations(47,48).

Importantly, although inter-patient variability remained the dominant source of overall proteomic diversity, protocol choice imposed a reproducible and biologically meaningful bias on tumor identity. These data indicate that dissociation should not be viewed as a neutral preparative step but rather as an intervention that selectively enriches specific cellular states while depleting others.

### Protocol-dependent preservation of tumor architecture and extracellular matrix

The protocol-dependent differences in cellular states described above were mirrored by striking differences in extracellular matrix (ECM) organization, highlighting the tight coupling between tissue architecture and tumor cell identity. A central finding of this ECM-focused analysis is that fragment-based CUT organoids preserve a markedly broader and more integrated repertoire of both core and associated matrisome proteins than dissociation-based DIS cultures. CUT organoids retained collagens, ECM glycoproteins, proteoglycans, ECM regulators, and secreted factors that assembled into highly interconnected protein networks linked to collagen fibrillogenesis, elastic fiber formation, integrin-mediated signaling, and vascular support.

These data indicate that mechanical fragmentation preserves endogenous, tumor-derived matrix architecture that is largely disrupted by enzymatic dissociation. In contrast, DIS organoids were dominated by basement membrane components associated with Matrigel exposure, as reflected by enrichment of laminins and type IV collagen together with the relative absence of higher-order ECM interaction networks. Because Matrigel is derived from murine Engelbreth–Holm–Swarm sarcoma, and many basement membrane proteins are highly conserved between mouse and human, species assignment of individual ECM peptides by standard human database searches remains challenging. We therefore interpret these signals conservatively, not as direct evidence of Matrigel carryover alone, but as an ECM state shaped by prolonged exposure to an exogenous basement membrane matrix. Similar matrix simplification after dissociation has been reported across tumor and developmental systems, where loss of interstitial ECM and altered mechanosignaling are accompanied by substitution with artificial matrices (8,10,49,50). This distinction may be particularly relevant in glioblastoma, where invasion and cellular plasticity are strongly regulated by matrix stiffness, topology, and integrin-dependent mechanotransduction(51–53). Preservation of native ECM architecture in CUT organoids may therefore better maintain biomechanical cues that influence tumor growth, migration, and tissue organization. Whether prolonged culture in defined exogenous matrices drives sustained transcriptional and proteomic reprogramming of tumor cells remains an important question for future study.

Importantly, although both protocols preserved a shared baseline matrisome, the selective depletion of interstitial and regulatory ECM components in DIS cultures fundamentally altered the physical and biochemical context experienced by tumor cells. This distinction is not merely structural. The ECM is now recognized as a dynamic regulator of tumor progression, influencing invasion, angiogenesis, lineage plasticity, immune interactions, and therapeutic response through both biochemical signaling and mechanical cues (54–57). By preserving endogenous ECM composition and organization, CUT organoids are therefore more likely to retain these microenvironmental inputs, whereas dissociation-based cultures represent a matrix-substituted and reductionist system.

Together, these findings identify ECM preservation as a major axis along which organoid culture strategies diverge. They further suggest that studies focused on ECM-dependent processes, including tumor–stroma interactions, invasion dynamics, mechanotransduction, and vascular niche signaling, are likely to be highly sensitive to protocol choice and would benefit from fragment-based approaches that preserve native matrix architecture.

### Implications for tumor modeling and translational relevance

A central criterion for evaluating tumor organoid models is the extent to which they faithfully recapitulate the molecular, cellular, and microenvironmental features of the parental tumor. By directly comparing organoids to matched primary tumor samples, we show that fragment-based CUT organoids consistently retain a closer proteomic resemblance to the original tissue than dissociation-based DIS cultures. This conclusion is supported across multiple orthogonal analyses, including dimensionality reduction, correlation metrics, protein overlap, pathway enrichment, and independent histological validation, indicating that the observed differences reflect biological fidelity, not technical artifacts.

Notably, CUT organoids shared a large cohort of enriched proteins with primary tumors, whereas DIS organoids retained only a minimal subset of primary tumor–specific features. These CUT-specific overlaps were enriched for pathways related to neural and glial identity, vascular and stromal components, and extracellular organization - hallmarks of glioblastoma complexity that extend beyond tumor-intrinsic signaling programs. Consistent with this, prior studies using fragment-based or minimally processed tumor models have shown that preservation of tissue architecture and stromal context enhances fidelity to the in vivo tumor state (2,58).

Importantly, these findings do not diminish the utility of dissociation-based organoid systems. DIS cultures offer clear advantages in technical reproducibility, scalability, genetic accessibility, and compatibility with high-throughput perturbation or screening platforms. They have also proven powerful for interrogating tumor cell–intrinsic signaling, drug sensitivity, and lineage plasticity under controlled conditions (1,7,48). However, our data indicate that dissociation and reaggregation impose a consistent directional bias toward growth-adapted cellular states and exogenously supported matrix environments - features that should be explicitly considered when interpreting experimental outcomes.

Together, these results suggest that the choice of organoid protocol should be guided by the biological question rather than convenience alone. Studies focused on tumor architecture, extracellular matrix– dependent signaling, cellular heterogeneity, or tumor–stroma interactions are likely best served by fragment-based approaches that preserve endogenous tissue context. In contrast, dissociation-based systems may be better suited to reductionist applications that prioritize uniformity, scalability, or genetic manipulation. More broadly, our work underscores that organoid culture protocols are not neutral technical choices but active determinants of biological state, with direct implications for translational relevance and cross-study comparability.

## Additional Information

## Acknowledgements

We acknowledge the core facility CELLIM of CEITEC, supported by the Czech-BioImaging large RI project (LM2023050 funded by MEYS CR), for their support with obtaining scientific data presented in this paper. CIISB, Instruct-CZ Centre of Instruct-ERIC EU consortium, funded by MEYS CR infrastructure project LM2023042 and European Regional Development Fund-Project "Innovation of Czech Infrastructure for Integrative Structural Biology” (No. CZ.02.01.01/00/23_015/0008175), is gratefully acknowledged for the financial support of the measurements at the CEITEC Proteomics Core Facility. Computational resources were provided by the e-INFRA CZ project (ID:90254), supported by MEYS CR. Finally, we thank Prof. Aleš Hampl for his support and valuable discussions.

## Authors’ contributions

JS conceived the study, designed and performed organoid culture experiments, analyzed data, and wrote the manuscript. OB conceived the study, performed proteomic data analysis, interpreted the data, and wrote the manuscript. KC contributed to experimental work and sample processing, wrote the manuscript, and prepared the final figures. MB and RJ provided patient material, clinical expertise, and contributed to study design and interpretation of the data. MH performed pathological evaluation and histological analyses. KK, MS, JO, and KAC contributed to experimental work, sample preparation, and technical support. ZH contributed to data interpretation, scientific discussion, and manuscript revision. DB conceived and supervised the study, secured funding, interpreted the data, and wrote the manuscript. All authors reviewed and approved the final manuscript.

### Ethics approval and consent to participate

Primary human tumor tissue was obtained from patients undergoing surgical resection after written informed consent. The study was approved by the Ethics Committee of St. Anne’s University Hospital Brno and the Faculty of Medicine, Masaryk University (reference number EK-FNUSA-27/2023). All procedures involving human participants were performed in accordance with institutional guidelines and with the ethical standards of the 1964 Declaration of Helsinki and its later amendments.

### Data availability

The mass spectrometry proteomics data generated in this study will be deposited in the ProteomeXchange Consortium via the PRIDE partner repository upon acceptance of the manuscript, and the accession number will be provided in the final published version. All other data supporting the findings of this study are available from the corresponding author upon reasonable request.

### Funding information

This research was supported by the Ministry of Health of the Czech Republic in cooperation with the Czech Health Research Council under project No. NW24-08-00157, by the Czech Science Foundation (Grant No. GA24-11357S), by Masaryk University (MUNI/A/1835/2025 and MUNI/A/1843/2025), and by the CREATIC project, funded by the European Union (Grant Agreement No. 101059788).

## Supplementary Information

**Supplementary Figure S1.**
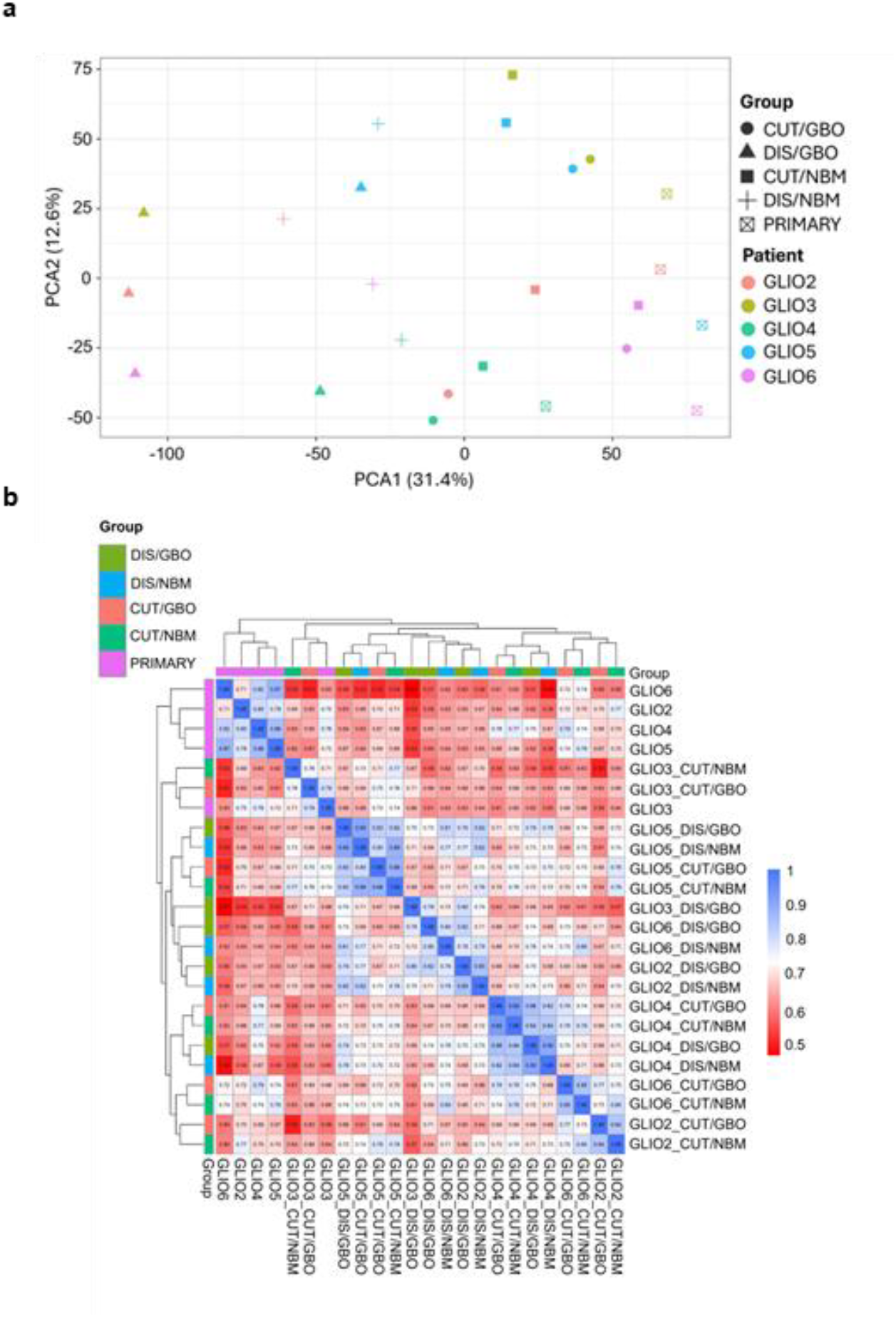
Assessment of biological variability and culture medium effects across all culture conditions. **(A)** PCA of proteomic profiles from all biological replicate samples, including CUT and DIS organoids cultured in GBO and NBM media, together with matched primary tumor samples. **(B)** Pearson’s correlation analysis of proteomic profiles across all biological replicate samples, illustrating relationships between culture conditions and primary tumors.

**Supplementary Table 1:** Clinical and pathological data of the patient samples used.

| Code | Clinical data |  |  |  |  | Pathology |  |  |  |  |  |  |
| --- | --- | --- | --- | --- | --- | --- | --- | --- | --- | --- | --- | --- |
|  | Gender | Age at the surgery | Localization | SVZ involvement | Epilepsy | Final integrated diagnosis according to the WHO CNS5 | IDH1 R132H IHC | IDH1/2 sequencing | pMGMT methylation | p53 IHC | ATRX IHC | Ki-67 |
| GLIO1 | Female | 54 | Left frontal lobe | NO | YES | Glioblastom IDH-wildtype, WHO G4 | wildtype | wildtype | unmethylated | mutant | retained | 30% |
| GLIO2 | Male | 51 | Left frontal lobe | YES | NO | Glioblastom IDH-wildtype, WHO G4 | wildtype | wildtype | unmethylated | wildtype | retained | 10% |
| GLIO3 | Male | 57 | Left frontal lobe | YES | NO | Glioblastom IDH-wildtype, WHO G4 | wildtype | wildtype | unmethylated | mutant | retained | 25% |
| GLIO4 | Male | 73 | Left temporo-parietal lobes | NO | NO | Glioblastom IDH-wildtype, WHO G4 | wildtype | wildtype | methylated | mutant | retained | 30% |
| GLIO5 | Male | 75 | Left temporal lobe | NO | NO | Glioblastom IDH-wildtype, WHO G4 | wildtype | wildtype | unmethylated | mutant | retained | 40% |
| GLIO6 | Female | 64 | Right temporo-parietal lobes | YES | NO | Glioblastom IDH-wildtype, WHO G4 | wildtype | wildtype | unmethylated | mutant | retained | 20% |

**Supplementary Table 2:** List of antibodies used for IHC.

| Antigen | Host/ antibody type | Clone | Working dilution | Manufacturer | Location |
| --- | --- | --- | --- | --- | --- |
| GFAP (glial fibrillary acidic protein) | Mouse monoclonal | 6F2 | 1:200 | Dako Agilent | Santa Clara, CA, USA |
| S100β | Rabbit polyclonal | Polyclonal | 1:1000 | Dako Agilent | Santa Clara, CA, USA |
| NSE (neuron-specific enolase) | Mouse monoclonal | MRQ-55 | Ready-to-use (RTU) | Cell Marque | Rocklin, CA, USA |
| CD31 | Mouse monoclonal | JC70A | 3:100 | Innovative Diagnostik-Systeme GmbH & Co. KG | Hamburg, Germany |

**Supplementary Table 3:** Processed proteomic data of technical replicates in an Excel spreadsheet

**Supplementary Table 4:** Processed proteomic data of biological replicates in an Excel spreadsheet

